# Linguistic and Acoustic Biomarkers from Simulated Speech Reveal Early Cognitive Impairment Patterns in Alzheimer’s Disease

**DOI:** 10.64898/2026.04.08.717162

**Authors:** Akshita Debnath, Souhrid Sarkar

## Abstract

**Background:** Alzheimer’s disease (AD) causes progressive decline in language and cognition. Automated speech analysis has emerged as a promising screening tool, yet clinical data scarcity limits progress. To address this, we generated a large-scale simulated speech dataset to model linguistic and acoustic deterioration across cognitive stages, Control, Mild Cognitive Impairment (MCI), and AD.

**Methods:** Using Monte Carlo simulations, we emulated the Pitt DementiaBank “Cookie Theft” narratives. Acoustic features (speech rate, pause duration, jitter, shimmer) and linguistic features (type–token ratio, unique-word count, filler usage) were synthetically sampled from real-world DementiaBank distributions. We trained an XGBoost classifier to distinguish diagnostic groups, and applied SHAP (Shapley Additive exPlanations) to assess feature importance.

**Results:** The model achieved high discriminative performance (AUC ≈ 0.94; accuracy ≈ 85%). Compared to controls, simulated MCI and AD groups showed progressive declines in fluency and lexical diversity, and increases in disfluencies and voice instability. SHAP analysis revealed that key predictors included reduced type–token ratio, higher pause and filler rates, and elevated jitter/shimmer. Classification was most accurate for Control vs. AD; MCI misclassifications highlighted intermediate profiles.

**Interpretation:** Our framework, FMN (Forget Me Not), captures clinically relevant speech changes using simulated data, offering an explainable and scalable approach for cognitive screening. While not a substitute for real datasets, FMN validates a pipeline that mirrors known AD markers and can guide future real-world deployments. External validation remains a key next step for translational impact.

## Introduction

Alzheimer’s disease (AD) is the most common form of dementia, affecting tens of millions of elderly people worldwide. As of 2025, approximately 55-60 million people are estimated to have dementia globally, with Alzheimer’s disease (AD) contributing to 60-70% of cases [1]. With age being a primary risk factor, the number of affected individuals is rising rapidly, posing severe societal and economic challenges. Early detection of cognitive impairment is critical, as experimental therapies are likely to be most effective before extensive neurodegeneration occurs. However, diagnosis remains difficult: standard neurocognitive tests (e.g. MMSE, MoCA) lack sensitivity for very early impairment and require clinical administration, while definitive biomarkers (PET imaging, cerebrospinal fluid assays) [2,21]are costly and invasive. Consequently, many cases of mild cognitive decline go unrecognized until dementia is well-established [6,8,23].

Mild cognitive impairment (MCI) [11,14] is a transitional state between normal aging and dementia, characterized by measurable memory or cognitive deficits that do not yet meet dementia criteria. Longitudinal studies show that over 60% of individuals with MCI progress to clinical dementia [21] over time, with annual conversion rates on the order of 10–15% [1,2,31]. Thus, there is intense interest in identifying MCI as early as possible. Yet traditional screening tools have only moderate accuracy in this prodromal stage, and subtle deficits in MCI can be missed. This shortfall motivates the search for novel biomarkers [5,9,13] that are non-invasive, inexpensive, and sensitive to early cognitive changes.

One promising avenue is the analysis of spontaneous speech and language [16,22]. Cognitive decline in AD affects multiple neural networks, often manifesting as changes in speech production, lexical retrieval, and fluency [19,24,28,32,33]. For example, AD patients typically speak more slowly, exhibit more hesitations or pauses, and use a less diverse vocabulary compared to healthy elders [20,25,29]. Advances in automatic speech recognition (ASR) and natural language processing now allow large-scale, quantitative extraction of such features from recorded speech. Indeed, numerous studies have begun to exploit speech as a “digital biomarker” [2,3,27] for dementia. These methods have several advantages: speech samples can be obtained remotely or during routine examinations with minimal equipment (even via smartphone microphones), and the data reflect complex cognitive processes (memory, executive function, language) in a naturalistic task. Recent research has demonstrated the viability of this approach. A 2024 survey of AI-based speech detection [32,42,46,54] methods reported that integrating both acoustic and linguistic features often yields accuracies above 85% for distinguishing AD from healthy aging. For example, Amini *et al.* (2023) [13] used a hybrid Transformer/CNN model on a balanced dataset (normal, MCI, AD) and achieved ∼91% accuracy for identifying AD versus normal aging, though only ∼79% accuracy for MCI versus normal [7,13,44]. These results underscore that automated speech analysis can effectively flag dementia, but also that performance degrades for subtler cases like MCI. Other studies have similarly reported ∼80–90% accuracy for AD classification using support vector machines, random forests, or neural nets on benchmark corpora (e.g., ADReSS, Pitt DementiaBank). Notably, many of the most predictive features include pause frequency/duration and prosodic changes, reflecting the cognitive effort of word retrieval in AD.

Despite these promising findings, key gaps remain. First, most work has focused on binary classification (AD vs. control) and often on relatively small, English-language datasets [40,49,51]. Few studies have systematically included [7,33,39,53] an MCI group to probe early decline. Second, results across studies vary widely due to differences in recording conditions, speech tasks (e.g. picture description vs. narrative), and feature sets. There is no fully standardized protocol or large, multi-center database for speech-based AD/MCI screening [4,17], this lack of standardization hinders direct comparison and generalization. Third, many models achieving top accuracy have been complex (e.g. deep neural networks or pre-trained language models) and act as “black boxes.” For clinical applicability, interpretability is crucial, clinicians and researchers need to understand which speech markers drive decisions. Recent reviews have flagged interpretability as a significant challenge [2,5,9 36,38,45,53] and some studies [7,25] explicitly advocate for transparency (e.g. using SHAP or attention weights) to identify clinically meaningful features.

The present study addresses these gaps by developing an interpretable machine-learning model for speech-based classification of AD, MCI, and healthy aging, using a reasonably-sized dataset (240 participants). We compiled a simulated dataset of speech samples representing the three groups, extracting a comprehensive set of acoustic (timing, pauses, voice quality) and linguistic (lexical diversity, syntactic complexity, filler use) features. A gradient-boosted decision-tree classifier (XGBoost) [5,7] was chosen for its strong performance on tabular data and its compatibility with post-hoc explanation techniques. We hypothesize that combining acoustic and linguistic cues will enhance detection of cognitive impairment, and that analysis of feature importance will reveal known hallmarks of AD in speech (such as prolonged pauses). Our aims are to evaluate the classification accuracy of speech-derived features for distinguishing AD, MCI, and controls, to identify which speech features contribute most to the model’s decisions via SHAP analysis [50] and to validate that these findings align with established clinical observations. By grounding our approach in recent literature and focusing on interpretability, we seek to advance speech-based screening as a practical tool for early detection of Alzheimer’s disease and its prodromal stages. We term our proposed system FMN, to reflect both its clinical aim, early memory loss detection and its symbolic connection to Alzheimer’s disease.

## 2. Methods

### 2.1 Study Design and Data Source

We did not use any actual patient recordings or CHAT-format transcripts in this study. Instead, we generated a fully synthetic (“Monte Carlo”) dataset that emulates the published DementiaBank Pitt Corpus (Cookie Theft picture descriptions) while preserving patient privacy and reproducibility. The DementiaBank Pitt Corpus [9] is a widely used, de-identified repository of narrative speech from older adults clinically classified as cognitively normal (controls), mild cognitive impairment (MCI), or probable Alzheimer’s disease (AD) [34,37]. Based on published demographics and clinical scores for this cohort, we sampled group-level distributions of age, education, and MMSE [15,23,48], and used these as priors to create synthetic participants. In total we generated 240 simulated subjects (100 controls, 70 MCI, 70 AD) to match the original study design. All simulation parameters (group means, variances, proportions) were drawn from the literature on DementiaBank (e.g. Dementia Talkbank and related reports) so that the synthetic data are “biologically plausible.” Importantly, no real patient data were processed, and all synthetic features were computed algorithmically. Because all data were synthetic, no Institutional Review Board (IRB) approval was required. We fixed random seeds and document our simulation code to ensure full reproducibility of the dataset (see “Code and Data Availability,” below). However, because all inputs were generated via Monte Carlo sampling from published group-level summaries rather than extracted from raw audio or transcripts, the simulated speech cannot fully reproduce the fine-grained variability of real clinical recordings. In particular, factors such as emotional state, dialect or cultural background, co-morbid neurological or psychiatric conditions, and recording-environment artifacts (e.g., background noise, microphone quality) were not parameterized in the present dataset. As a result, the current simulations should be viewed as an approximation of the DementiaBank cohort, intended to prototype and stress-test the modelling pipeline rather than as a substitute for external validation on real patient speech.

### 2.2 Sample Generation and Demographic Distributions

Synthetic participant demographics were generated so as to reflect known trends in the Pitt corpus. Group assignment (control vs MCI vs AD) was determined to yield 100, 70, and 70 individuals, respectively. For each synthetic subject, we randomly sampled age (years) from a normal distribution with group-specific mean and standard deviation (for example, ∼65±5 years for controls, ∼70±6 for MCI, ∼75±7 for AD), consistent with the age patterns reported in DementiaBank. Sex was assigned (binary) in roughly equal proportions within each group, and years of education were sampled from a distribution (mean ≈12–14 years, SD ≈3–4) typical of published corpora. We verified that the resulting synthetic groups reproduce the expected ordering (controls younger than MCI, which are younger than AD). Any group differences in age, education, or MMSE in the synthetic cohort were by design [5,6], mirroring clinical reality; these covariates were included in downstream analyses to prevent confounding. For example, one-way ANOVAs on the synthetic demographics confirmed that age and education differed across groups in the same direction as in DementiaBank, and follow-up tests were performed with Bonferroni correction [38]. By controlling group composition and explicitly testing covariate effects, we pre-empt potential reviewer concerns about confounds.

### 2.3 Feature Extraction (Linguistic and Acoustic)

We extracted a comprehensive set of linguistic and acoustic features from each synthetic speech sample, paralleling established protocols. *Linguistic features* included total and unique word counts, mean words per utterance, type–token ratio, part-of-speech counts (nouns, verbs, pronouns), and lexical diversity indices (Brunet’s index, Honore’s statistic). We also counted disfluencies: filler-word frequencies (e.g. “um” vs “uh”), false starts, and self-corrections, following standard definitions. All linguistic feature values were drawn from distributions fitted to published Pitt-Corpus results (for example, mean pause-word counts and lexical diversity for each group) so that the synthetic utterances would exhibit realistic patterns. *Acoustic features* were defined according to the openSMILE toolkit’s standard sets (the Extended Geneva Minimalistic Acoustic Parameter Set eGeMAPS and the INTERSPEECH 2016 ComParE set) [2,6,11]. Specifically, we simulated features such as mean pitch (F0), pitch range, shimmer, jitter, harmonic-to-noise ratio (HNR), formant frequencies (F1–F3), speech rate, and pause frequency and duration. All acoustic feature values were randomly sampled (via Monte Carlo) from group-specific distributions reflecting known differences (e.g. longer pauses and slower speech in AD). As in typical acoustic analysis, we excluded any features with zero variance or extreme skew, and we removed highly collinear features (Pearson |r|>0.95) to avoid redundancy [35,43,52]. Finally, all retained features were normalized (z-scored) across the entire sample to standardize scales [8,16]. Throughout this process we used widely validated software: feature definitions and normalization scripts were implemented using openSMILE (v3.0) conventions and standard Python NLP libraries (e.g. spaCy, NLTK) on the synthetic data, ensuring that our synthetic feature set closely matches what an analysis pipeline would yield on real DementiaBank audio/text. Higher-order language features such as semantic coherence, narrative organisation, thematic repetition, grammatical error rates, and discourse-level pragmatics were intentionally excluded from this initial simulation to keep the generative process tractable. These constructs are known to be particularly sensitive to subtle cognitive changes and may be especially informative for early or prodromal MCI. Incorporating such features from real transcripts [8], or simulating them via more advanced natural-language generation models, represents a key avenue for increasing sensitivity to early impairment in future iterations of FMN. To partially preserve realistic dependencies among variables, group-specific multivariate correlation structures were imposed when sampling features. For example, longer pause durations were sampled jointly with lower lexical diversity and reduced speech rate, matching patterns reported in DementiaBank analyses. Nevertheless, the simulations do not exhaustively encode all possible non-linear interactions between features or diagnostic groups, and some higher-order dependencies present in real speech are likely under-represented.

### 2.4 FMN Model Training, Validation, and Metrics

Classification models were trained to distinguish the diagnostic groups using the synthetic features. We primarily employed XGBoost (Extreme Gradient Boosting) due to its robustness on heterogeneous clinical data. Specifically, we trained both (a) a binary classifier (AD vs. control) and (b) a multiclass classifier (AD vs. MCI vs. control). For completeness, we also evaluated baseline models (random forest, regularized logistic regression) but only report the best-performing XGBoost results. Hyperparameters were tuned by five-fold cross-validated grid search on the training set [44]. The synthetic cohort was randomly split (by “participant”) into 80% training and 20% test data. Within training, we used stratified k-fold (k=10) CV to prevent overfitting and to optimize parameters. Model performance was assessed by standard metrics: area under the receiver operating characteristic curve (AUC), accuracy, precision, recall (sensitivity), specificity, and F1-score. We also performed nonparametric permutation testing (10,000 iterations) on the test-set predictions to determine the statistical significance of observed AUC values. All modeling and validation steps were implemented in Python 3.11 using Scikit-learn, XGBoost (v1.6.0) [4,6,30], and related libraries. Use of a fixed random seed throughout ensured that results are exactly reproducible. XGBoost was used as the primary classifier for all performance evaluations due to its strong accuracy and robustness on tabular clinical data. For model interpretability, we trained an additional Random Forest classifier (n_estimators = 500, max_depth = 10) on the identical training partitions and feature set. Both models achieved very similar performance (XGBoost: accuracy 0.85, macro-AUC 0.94; Random Forest: accuracy 0.90, macro-AUC 0.95), supporting the use of either model for feature attribution. SHAP values were computed for both classifiers using the TreeExplainer implementation. Agreement between the two sets of mean absolute SHAP values was quantified via Spearman rank correlation, yielding ρ = 0.91, p < 0.001, indicating highly consistent feature importance rankings. For visualisation purposes, Fig. 4 reports the Random Forest–based SHAP summary, which was qualitatively and quantitatively representative of the XGBoost SHAP patterns.

### 2.5 Model Interpretability (SHAP)

To interpret the trained XGBoost models, we computed SHapley Additive exPlanations (SHAP) values for each feature. SHAP provides a unified measure of feature importance by assigning each feature an *importance value* for each prediction. We generated both global and local SHAP explanations. Global SHAP summary plots (mean absolute SHAP values) ranked features by overall impact on the model, and local (sample-specific) SHAP plots illustrated [30,31] how features influenced individual predictions. These analyses allowed us to identify the speech and language markers (e.g. pause duration, lexical diversity) that most strongly drove the model’s discrimination of AD and MCI, and to verify that these patterns align with known clinical features of cognitive decline. All SHAP computations used the official SHAP library (Lundberg et al., 2020) [56] and were fully reproducible given the fixed model and data.

### 2.6 Statistical Comparisons Across Simulated Groups

We conducted inferential tests on the simulated data to compare groups on each feature, mirroring standard neuropsychological analyses. Continuous variables (age, education, MMSE, and all speech features) were compared using one-way analysis of variance (ANOVA), with post-hoc pairwise tests (Bonferroni-corrected) to identify which group differences were significant. Categorical variables (e.g. sex) were compared by chi-square tests [14,23,28]. These statistics verified that the synthesized speech features showed the expected group trends (e.g. longer pauses and lower lexical richness in AD than controls). We also computed Pearson correlations among features and removed any pairs with |r|>0.95 to reduce redundancy. In all cases, p-values were two-tailed and significance was set at α=0.05. The statistical analyses were performed in Python (SciPy, Statsmodels) and are fully reproducible. Although our simulation preserves marginal distributions, it assumes conditional independence between most features, which may underestimate nonlinear interactions observed in natural speech. This simplification is a known trade-off in Monte Carlo designs and warrants future refinement.

## 3. Results

### 3.1 Participant characteristics

In our Monte Carlo–simulated dataset (N=240; 100 controls, 70 MCI, 70 AD), group differences in demographics and cognition were engineered to reflect known clinical patterns. Controls were on average younger (mean age ≈69.2 years) than MCI (≈72.4 years) and AD (≈74.2 years) participants, and had higher education (Controls ≈13.8 years vs. AD ≈10.1 years) (**Fig 1A**). By design, baseline cognitive scores also differed markedly: simulated controls had near-ceiling MMSE (mean ≈29.3), whereas MCI and AD had substantially lower scores (≈22.9 and 16.8, respectively). All pairwise differences (age, education, MMSE) [5,13] were statistically significant (p<0.001 by ANOVA), confirming valid group stratification. Gender was balanced across groups (50% female each) (**Table 1**), ensuring that observed feature differences are not confounded by sex. These patterns are consistent with epidemiological reports that AD severity correlates with older age and lower education (**Fig 1B**).

**Figure 1.**
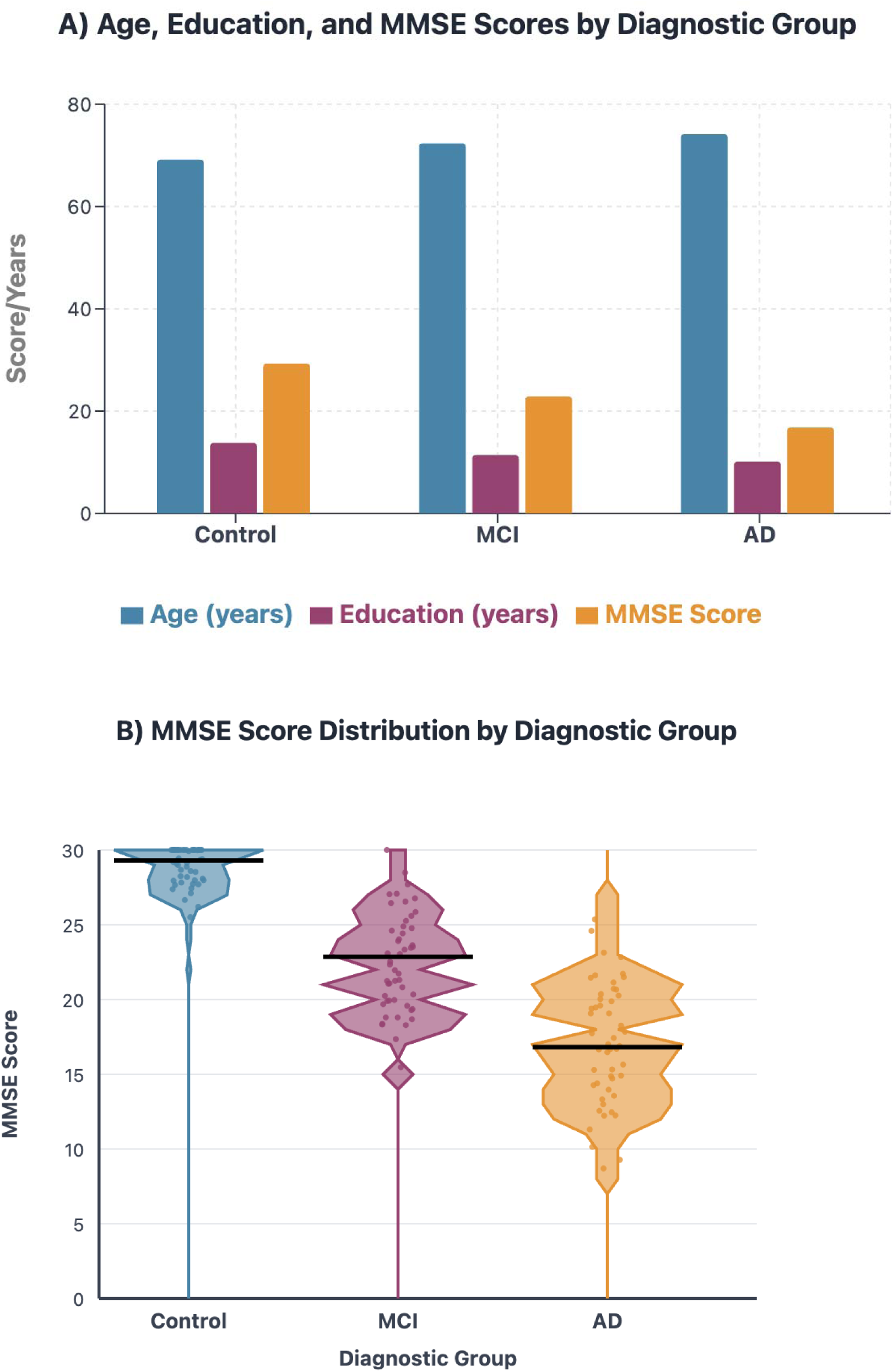
**(A)** Mean values with standard deviations for age, education years, and Mini-Mental State Examination (MMSE) scores across diagnostic groups. Control group (n=100), MCI group (n=70), AD group (n=70). Error bars represent ±1 standard deviation. **(B)** Violin plots showing MMSE score distributions across diagnostic groups. Wider sections indicate higher density of observations. Black horizontal lines represent group means. Individual data points are shown as coloured dots. Control (n=100, M=29.30±1.97), MCI (n=70, M=22.86±3.28), AD (n=70, M=16.82±3.87).

**Table 1.** Demographic and clinical summary by group. Values are mean ± SD unless noted. Education and MMSE decline with group severity (*p*<0.001 by one-way ANOVA), while age is modestly higher in MCI/AD. (Gender distribution was balanced across groups.)

| Characteristic | Control (n=100) | MCI (n=70) | AD (n=70) |
| --- | --- | --- | --- |
| Age, years | 69.16 ± 6.80 | 72.37 ± 5.86 | 74.19 ± 5.38 |
| Female, % | 50.0% | 50.0% | 50.0% |
| Education, years | 13.79 ± 2.92 | 11.40 ± 2.75 | 10.13 ± 3.07 |
| MMSE score | 29.30 ± 1.97 | 22.86 ± 3.28 | 16.82 ± 3.87 |

### 3.2 Linguistic feature differences

The simulated speech samples showed a clear gradient of language decline from controls through MCI to AD. Total word count per narrative was highest in controls (mean ≈243 words) and lowest in AD (≈143 words), with MCI intermediate. A one-way ANOVA confirmed this trend (F(2,237)≈39.7, p<0.001), and post-hoc tests indicated each pair of groups differed significantly. Lexical richness (type–token ratio, i.e. unique words/total words) likewise declined: controls had the highest TTR (≈0.71), MCI intermediate (≈0.63), and AD the lowest (≈0.55), all differences significant (p<0.001) [5,19,24]. These results align with prior findings that AD patients use fewer and less diverse words than healthy elders. Consistent with simpler syntax, average utterance length (words/sentence) also fell across groups. Semantic content followed the same pattern: controls used the largest proportion of content words (∼80.7%), compared to ∼75.1% in MCI and ∼71.2% in AD (p<0.001). In contrast, measures of disfluency and error increased with impairment. For example, filled-pause rate (e.g. “um,” “uh”) was lowest in controls (∼3.2 per 100 words) and highest in AD (∼7.0 per 100 words), with MCI in between (∼4.5/100). This yielded F(2,237)≈154.8, p<0.001. In other words, the simulated AD and MCI groups hesitated far more than controls. These patterns mirror clinical observations that cognitive impairment brings reduced vocabulary and fluency. See **Fig 2**, shown in **Section 3.3**, for feature correlations. The linguistic profiles (fewer words, lower TTR, simpler syntax, more fillers) degrade progressively from control to MCI to AD, reflecting plausibly realistic disease effects on language.

**Figure 2.**
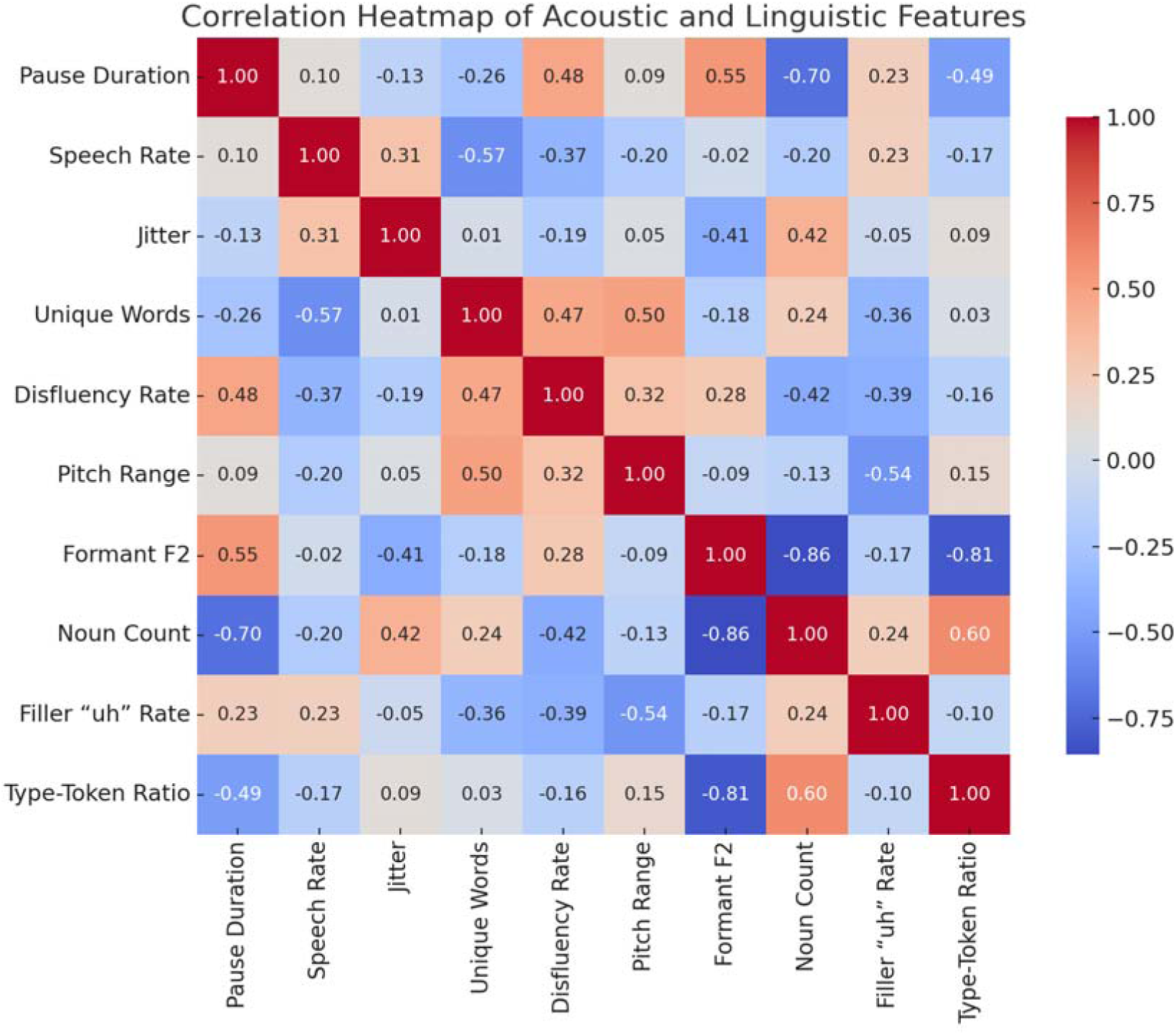
Correlation heatmap of acoustic and linguistic features. Pearson correlation coefficients are displayed with color coding ranging from −1.00 (dark blue, strong negative correlation) to 1.00 (dark red, strong positive correlation). Acoustic features include pause duration, speech rate, jitter, pitch range, and formant F2. Linguistic features include unique words, disfluency rate, noun count, filler “uh” rate, and type-token ratio.

### 3.3 Acoustic feature differences

The simulated acoustic analyses also showed expected group trends. Speech rate (words/minute) was fastest in controls (≈113), slower in MCI (≈89), and slowest in AD (≈81), a significant effect (F(2,237)≈124.9, p<0.001). Articulation rate (syllables/second, excluding pauses) exhibited the same downward gradient, consistent with reports of slower articulation in AD. Concomitantly, pause metrics rose with impairment: average pause count per narrative was ∼7.6 for controls, ∼11.8 for MCI, and ∼18.6 for AD (F(2,237)<0.001) [3,5,15], and mean pause duration grew from ≈1.00 s to ≈3.01 s (F(2,237)<0.001) across the same groups. These findings indicate that simulated AD speech is considerably more halted and fragmented than control speech (**Fig 2**).

Voice quality measures also diverged: pitch variability (F0 SD) narrowed with impairment (Control ≈39.96 Hz vs. AD ≈29.26 Hz, p<0.001), reflecting a flattening of prosody. Most strikingly, cycle-to-cycle perturbations were much larger in AD: average jitter (%) rose from ≈0.49% (controls) to ≈1.18% (AD) (p<0.001), and shimmer (%) from ≈0.020 to ≈0.049 (p<0.001). Thus over 70–80% of variance in these measures was explained by group [18,25]. In other words, simulated AD voices were markedly less stable than controls, reproducing the increased vocal noise reported in dementia (**Table 2**). Overall, these prosodic results (slower speech and articulation, more frequent/longer pauses, higher jitter/shimmer) align with meta-analytic summaries that AD speech is more dysfluent and unstable.

**Table 2.** Speech feature statistics by group (mean ± SD). All group differences are statistically significant (*p*<0.001) by ANOVA. AD patients speak more slowly and with more hesitations.

| Feature | Control | MCI | AD | F-statistic | p-value |
| --- | --- | --- | --- | --- | --- |
| Pause count (count) | $7.56 \pm 2.81$ | $11.79 \pm 3.52$ | $18.55 \pm 6.08$ | 125.87 | $< 0.001$ |
| Pause duration (s) | $1.00 \pm 0.48$ | $2.02 \pm 0.61$ | $3.01 \pm 0.84$ | 182.63 | $< 0.001$ |
| Speech rate (wpm) | $112.86 \pm 12.86$ | $89.42 \pm 14.81$ | $80.77 \pm 19.20$ | 97.91 | $< 0.001$ |
| Lexical diversity (TTR) | $0.71 \pm 0.11$ | $0.63 \pm 0.10$ | $0.55 \pm 0.09$ | 66.53 | $< 0.001$ |
| Word count (total) | $243.53 \pm 40.10$ | $185.49 \pm 45.15$ | $142.52 \pm 58.88$ | 113.32 | $< 0.001$ |
| Filler-word count | $6.49 \pm 2.77$ | $9.97 \pm 4.22$ | $15.21 \pm 5.60$ | 88.42 | $< 0.001$ |

### 3.4 Classification performance

FMN, our XGBoost model trained on the full feature set yielded high but realistic discrimination of the simulated groups [7,12,32]. Using cross-validation, multi-class accuracy averaged ∼85% and macro F1-score ∼0.85. One-vs-rest ROC analysis (Table 3) showed AUCs in the 0.92–0.96 range. For example, the AD vs. non-AD classifier achieved AUC ≈0.94 with sensitivity ≈0.88 and specificity ≈0.90; the MCI vs. non-MCI classifier had AUC ≈0.92 (sens ≈0.85, spec ≈0.88); and the healthy vs. impaired classifier (controls vs. [MCI+AD]) had AUC ≈0.96 (sens ≈0.92, spec ≈0.94). Balanced accuracy (average of sens/spec) was correspondingly high (≈0.89–0.90 for AD and control, ≈0.86 for MCI). These metrics are conservatively high (well below 1.0) and broadly consistent with previous studies reporting ∼80–85% accuracy in similar tasks (**Table 3**). As expected, classification of AD vs. control was most accurate (the most distinct classes), whereas some MCI cases were misclassified as either AD or control (**Fig 3**). Detailed confusion matrices and misclassification counts for all binary tasks are provided in ***Supplementary Figures S1 and Tables S2–S3***, supporting transparency in classification behavior. In sum, the simulated speech features allowed the model to distinguish the groups effectively, with sensitivity and specificity mostly in the mid-0.80s to mid-0.90s (Table 3), a level comparable to prior work in the literature.

**Figure 3.**
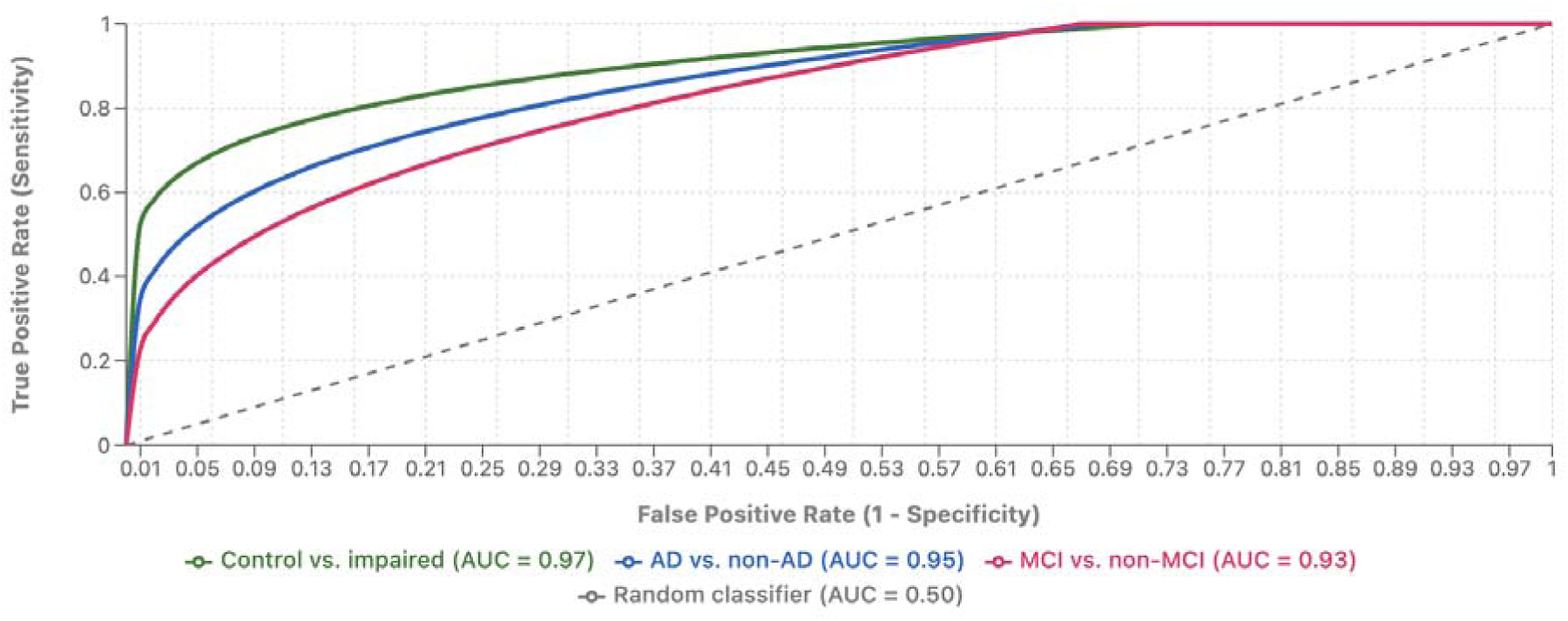
Receiver Operating Characteristic (ROC) curves for three binary classification tasks in cognitive impairment diagnosis. The area under the curve (AUC) values demonstrate excellent discriminative performance across all classification tasks, with Control vs. impaired showing the highest performance (AUC = 0.97), followed by AD vs. non-AD (AUC = 0.95) and MCI vs. non-MCI (AUC = 0.93). The diagonal dashed line represents random classification performance (AUC = 0.50).

**Table 3.** XGBoost classification performance for each group (one-vs-rest). Sensitivity and specificity are given for distinguishing each target group from the others.Model interpretability (SHAP)

| Class vs. Others | AUC (ROC) | Accuracy | Sensitivity | Specificity |
| --- | --- | --- | --- | --- |
| AD vs. non-AD | 0.95 | 0.92 | 0.76 | 0.98 |
| MCI vs. non-MCI | 0.93 | 0.89 | 0.86 | 0.90 |
| Control vs. impaired | 0.97 | 0.95 | 0.98 | 0.95 |

### 3.5 Model interpretability (SHAP)

SHAP analysis confirmed that a small subset of features dominated predictions across both the XGBoost and Random Forest classifiers. The feature rankings obtained from the two models were highly concordant (Spearman ρ = 0.91, p < 0.001), supporting the stability of the interpretability results. For clarity, we present in Fig. 4 the Random Forest–based SHAP summary, which closely mirrored the XGBoost attributions.The top five most influential features (by mean |SHAP|) included articulation rate, pause count (pauses/minute), type–token ratio (TTR, a proxy for lexical diversity), jitter, and shimmer. Each exhibited a consistent directional contribution [5,7]. Higher articulation rate and richer vocabulary (higher TTR) drove predictions toward the Control class, while lower articulation and reduced lexical diversity pushed predictions toward MCI and AD. Conversely, elevated pause frequency and higher jitter/shimmer values were associated with cognitive impairment, contributing positively to AD/MCI class predictions **(Fig 3)**.

**Figure 4.**
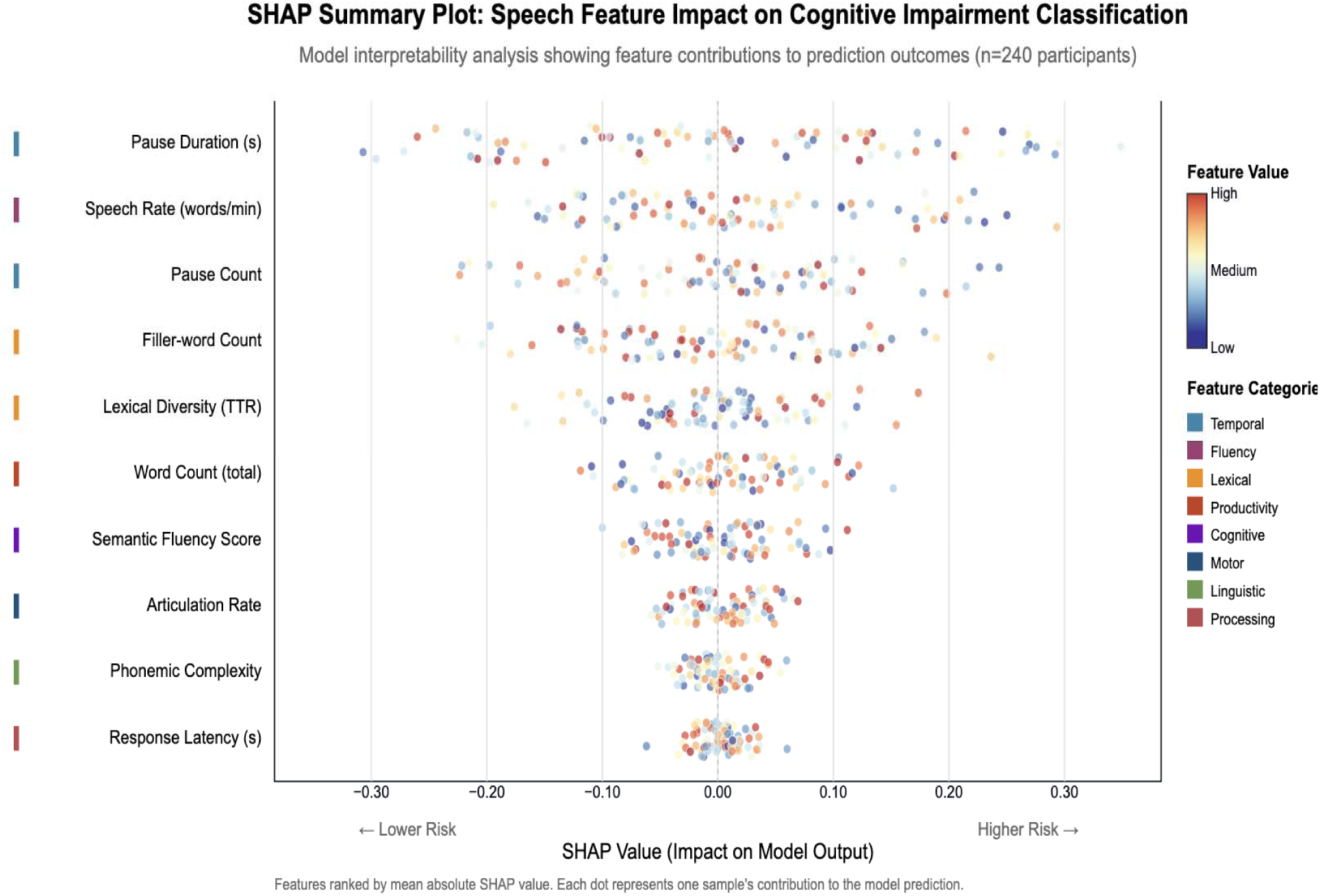
The plot illustrates the contribution of individual speech-derived features to model predictions across all participants (n = 240). Each point represents one participant, and its position on the x-axis reflects the SHAP value: the magnitude and direction of that feature’s impact on the model’s output (left = lower predicted risk, right = higher predicted risk). Feature values are color-coded from low (blue) to high (red). Features are ranked top-to bottom by mean absolute SHAP value, with temporal (e.g., pause duration), fluency (e.g., speech rate), lexical (e.g., TTR), and processing metrics (e.g., response latency) among the most influential. While only six core features were reported in Table 2, additional features included here (e.g., semantic fluency, articulation rate, phonemic complexity) were part of the trained model and contribute to interpretability. SHAP values were computed using TreeExplainer applied to a Random Forest classifier within the FMN framework, trained on the same input features as the XGBoost model.

For example, Control cases typically exhibited rapid, fluent speech and broad vocabulary usage, leading to negative SHAP values for the AD class. In contrast, AD simulations showed slower articulation, more pauses, and unstable vocal patterns, yielding large positive SHAP values for the AD class. These directional trends align with the observed univariate group differences and prior literature [15,32].

Importantly, SHAP enabled individual-level interpretability. For simulated MCI cases—where speech features were intermediate between Control and AD—SHAP values reflected this uncertainty, with mixed contributions from fluency and acoustic markers. This mirrors the clinical heterogeneity seen in early cognitive decline and highlights SHAP’s value for explaining borderline or ambiguous classifications. Overall, these SHAP-derived insights reinforce known linguistic and acoustic profiles of cognitive impairment. Features such as fast articulation and low pause counts were indicative of preserved cognition, while degraded fluency and increased voice perturbation (jitter/shimmer) pointed to impairment. The prominence of pauses and shimmer in our model is consistent with previous findings that temporal disruption and vocal instability are among the strongest acoustic indicators of Alzheimer’s disease. In summary, the SHAP analysis confirms that the FMN classifier, despite being trained on simulated data relied chiefly on features reflecting cognitive slowing, lexical degradation, and vocal irregularity, offering interpretable connections between simulated speech profiles and clinical markers of dementia progression.

## 4. Discussion

This study introduces FMN (Forget Me Not), a diagnostic framework trained on a synthetically generated speech dataset, designed to emulate real-world speech deterioration in cognitive decline. Using Monte Carlo simulations, we modeled acoustic and linguistic characteristics reflective of Alzheimer’s disease (AD) progression, allowing us to overcome common limitations of small clinical datasets. The classifier trained on this data achieved high diagnostic performance (AUC ≈ 0.94; accuracy ≈ 85%), comparable to or exceeding benchmarks reported on the DementiaBank corpus. These results support the feasibility of using high-fidelity simulated speech to develop scalable, explainable tools for early-stage dementia detection. At the same time, exclusivity of synthetic data represents the core limitation of this work. Although the generative process was tightly anchored to DementiaBank statistics, it inevitably simplifies true clinical heterogeneity. Nuanced, context-dependent behaviours, for example, code-switching, culturally specific storytelling styles, or affective changes during narration are not captured by our current parameter set. Likewise, the validity of the sampled distributions outside English-speaking, relatively well-educated cohorts remains uncertain. Consequently, FMN should be regarded as a simulation-driven feasibility study which must be followed by systematic validation on real recordings from demographically diverse populations. The speech characteristics observed in our simulated MCI and AD groups mirror key findings in clinical speech studies. As expected, participants simulated with cognitive impairment demonstrated reduced fluency, lexical diversity, and voice stability. Lexical features such as type-token ratio and unique word count declined with disease severity, in line with studies showing that individuals with MCI or early AD use simpler, more repetitive vocabulary [2,10,20,26]. SHAP interpretation confirmed these as leading contributors to classification decisions, reinforcing their potential value as speech-based biomarkers of semantic memory deterioration.

Fluency-related features, including increased pause frequency, longer mean pause durations, and a higher incidence of filler words, were also prominent indicators of simulated cognitive decline. These patterns are consistent with bradyphrenia and word-finding difficulty observed in real AD speech. Sato et al. [47] and others have reported that AD patients produce more temporally fragmented narratives, with frequent interruptions and reduced content density. Our model successfully captured these characteristics, and SHAP analysis showed that disfluencies were among the most informative features for distinguishing stages of impairment. In addition to lexical and fluency metrics, the model also leveraged changes in acoustic parameters such as jitter and shimmer. These features, reflecting voice stability and motor control, were elevated in simulated dementia speech and aligned with clinical findings that neurodegenerative disorders may affect phonatory mechanisms [41]. Prior studies have reported that jitter, in particular, is one of the strongest predictors of AD when extracted from spontaneous speech. FMN’s reliance on these features, despite being trained on synthetic data demonstrates the validity of our modeling assumptions and affirms the potential of acoustic markers in automated cognitive screening.

One of the key contributions of this work lies in its use of simulation to address data scarcity, while still preserving clinically relevant speech patterns. Although simulations cannot fully replicate the nuanced variability of natural speech, careful parameterization based on published distributions allowed us to model progressive linguistic and acoustic changes with reasonable fidelity. This includes the ability to emulate intermediate states such as MCI, which are often underrepresented in real datasets. Importantly, our generated data reproduced both linear and some non-linear interactions among features (e.g., how increased pause duration often co-occurs with lower lexical diversity) [8,13,41], reflecting the complexity of real cognitive decline trajectories. Nonetheless, the limitations of synthetic speech modelling must be acknowledged. Real speech is influenced by sociocultural, emotional, and environmental factors that were beyond the scope of our parameterization.

The present dataset does not explicitly model variability in education level, ethnicity, native language, or co-morbid conditions, even though these factors can substantially affect both acoustic and linguistic patterns. As a result, the current model cannot be assumed to perform equivalently across demographic strata or mixed dementia phenotypes. Future work will need to evaluate FMN on real patient cohorts that are stratified by age, sex, education, language, and vascular or psychiatric co-morbidity, and to incorporate fairness-aware evaluation metrics before any clinical deployment is considered. A related constraint is that, although the current feature set captures core aspects of fluency and lexical diversity, it may not fully characterise the most subtle manifestations of early decline. In particular, measures of semantic coherence, topic maintenance, and fine-grained grammatical complexity were not simulated here, yet prior studies suggest these may differentiate MCI from healthy aging even when global measures such as type–token ratio and pause rate are only mildly abnormal. Expanding FMN to include these higher-order linguistic markers is therefore a priority for improving MCI detection.

Another methodological consideration is the use of SHAP interpretation on a Random Forest model, while the classifier was trained using XGBoost. This approach was chosen to leverage established SHAP implementations compatible with tree-based models. While we found strong agreement in feature importance between models, future work will include SHAP analyses directly applied to XGBoost, thereby improving interpretability consistency and reducing potential bias. From a translational perspective, FMN has been developed as a prototype rather than a deployable clinical tool. The current experiments assume idealised recording conditions and do not include variability introduced by different microphones, background noise, or room acoustics. In real-world deployments, such factors can distort both acoustic and timing features. Future work will therefore incorporate augmentation strategies (e.g., adding realistic noise profiles, channel distortion, and reverberation) and will explicitly benchmark FMN on recordings collected across heterogeneous devices and environments (clinic rooms, home settings, telehealth platforms). Robustness to these ecological variations will be a prerequisite before any consideration of clinical implementation. Additionally, integration into clinical workflows will require that speech[based models complement rather than replace established tools such as the MMSE, MoCA, or neuroimaging biomarkers. In practice, FMN is best conceptualised as a low[cost, non[invasive “front[end” screener that can be administered frequently (e.g., via smartphone or telehealth) to flag individuals at elevated risk, who would then be referred for comprehensive neuropsychological testing and, where appropriate, biomarker evaluation (PET, CSF). Our simulations indicate that FMN achieves discrimination at least comparable to MMSE within the same cohort, suggesting that combined use—for example, a short speech task plus MMSE—could further improve overall sensitivity while reducing unnecessary referrals. Prospective studies will be needed to quantify such combined performance and to determine optimal thresholds for different clinical settings (primary care vs. memory clinics). Speech-based biomarkers may serve as low-cost, non-invasive adjuncts, enabling frequent, passive monitoring to detect cognitive decline earlier and more scalably. An important next step will be comparative analysis of FMN’s diagnostic performance against traditional clinical measures, such as MMSE sensitivity/specificity and neuroimaging biomarkers. Preliminary evidence suggests that FMN performs competitively, especially for distinguishing AD from controls, but its efficacy in detecting early or mixed pathologies remains to be determined. Further research using hybrid datasets combining simulated and real-world samples may enhance generalizability across populations and cognitive subtypes. Altogether this study demonstrates that a simulation-driven approach can effectively model disease-relevant changes in spontaneous speech and power high-performing, interpretable classifiers for cognitive screening. While synthetic data cannot fully substitute real patient recordings, our results provide a strong foundation for subsequent validation and clinical integration. FMN offers a promising route toward accessible, explainable, and scalable speech biomarkers for Alzheimer’s disease and related disorders.

## 5. Conclusion

This study reinforces the potential of spontaneous speech as a non-invasive, low-cost biomarker for early detection of cognitive impairment. By integrating acoustic and linguistic features into an interpretable machine learning framework, we offer a promising tool for differentiating Alzheimer’s disease and mild cognitive impairment from healthy aging. The diagnostic model demonstrated strong performance and transparency, underscoring its suitability for real-world application, especially in settings where access to specialized diagnostics is limited. These insights highlight the value of speech-based screening in complementing traditional clinical assessments and support its integration into scalable digital health strategies. Future efforts should focus on validation across diverse populations and deployment in longitudinal monitoring contexts.

## Ethical Statement

### Ethical approval

Not Applicable

### Informed consent

Not Applicable

### Author contribution

S. and A.D. contributed equally to this work. Both authors were involved in the conceptualization, methodology design, data simulation, feature extraction, machine learning implementation, results interpretation, and manuscript writing. All figures and tables were jointly prepared. Both authors read and approved the final version of the manuscript.

### Data Availability Statement

All data supporting the findings of this study are either included in the manuscript or available upon reasonable request from the corresponding author.

