## Supplementary figures and images for "Linguistic and Acoustic Biomarkers from Simulated Speech Reveal Early Cognitive Impairment Patterns in Alzheimer’s Disease"

### Supplemental Data 1

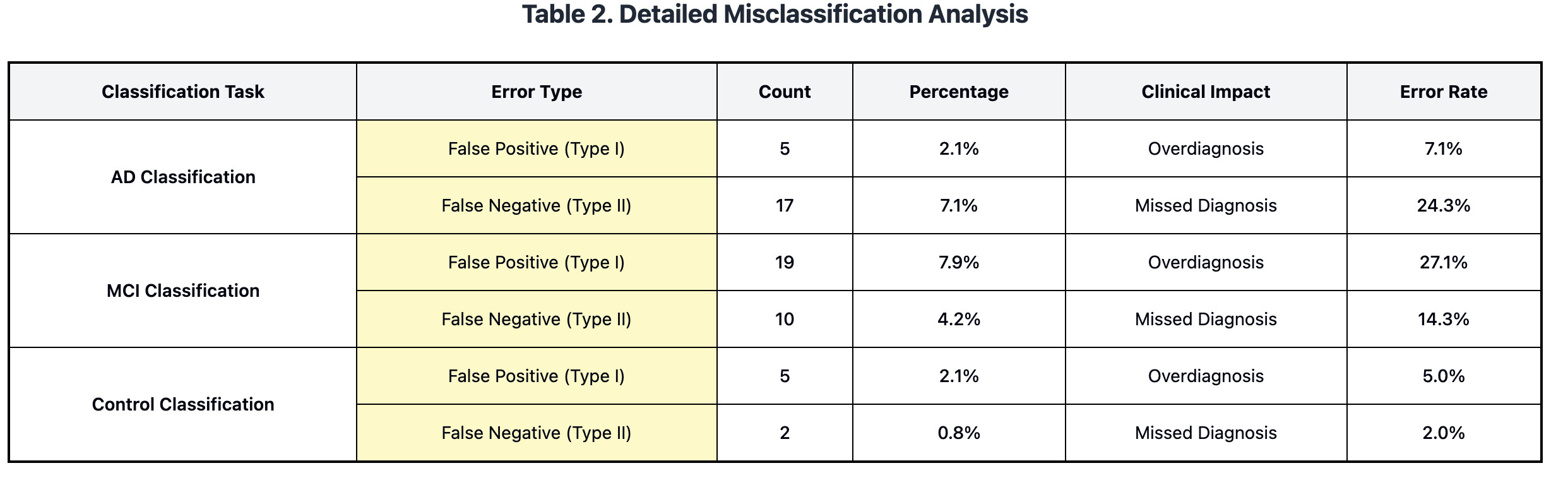

### Supplemental Data 2

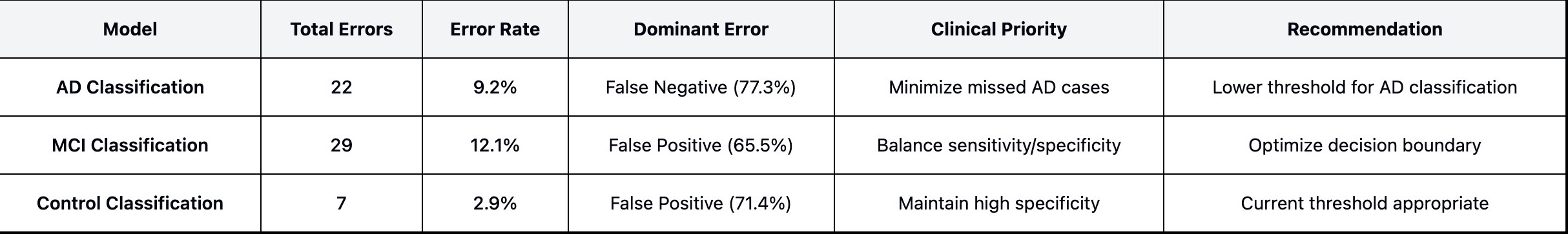

### Supplementary Figures S1

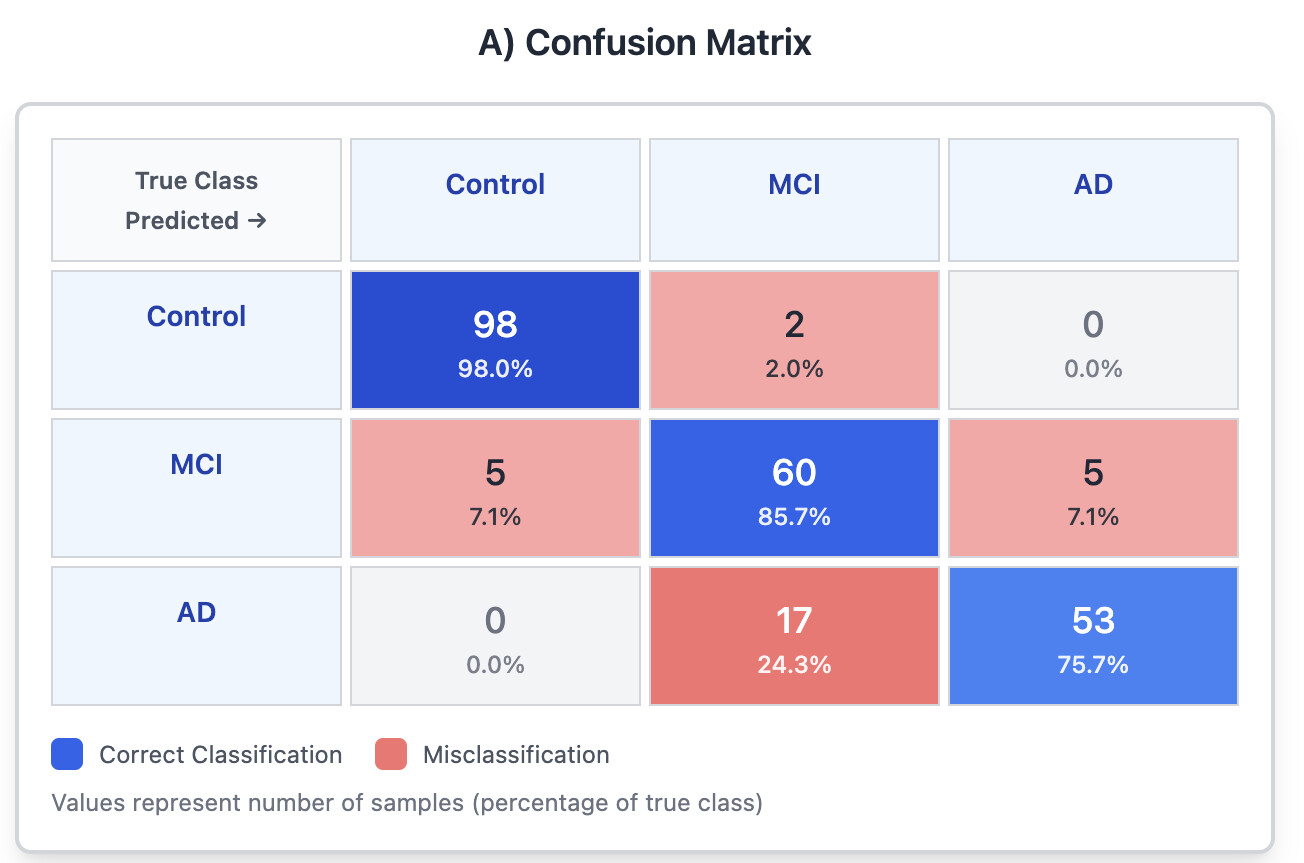
