## Supplementary Table S1 for "Linguistic and Acoustic Biomarkers from Simulated Speech Reveal Early Cognitive Impairment Patterns in Alzheimer’s Disease"

| Classification Task | Error Type | Count | Percentage | Clinical Impact | Error Rate |
| --- | --- | --- | --- | --- | --- |
| AD Classification | False Positive (Type I) | 5 | 2.1% | Overdiagnosis | 7.1% |
|  | False Negative (Type II) | 17 | 7.1% | Missed Diagnosis | 24.3% |
| MCI Classification | False Positive (Type I) | 19 | 7.9% | Overdiagnosis | 27.1% |
|  | False Negative (Type II) | 10 | 4.2% | Missed Diagnosis | 14.3% |
| Control Classification | False Positive (Type I) | 5 | 2.1% | Overdiagnosis | 5.0% |
|  | False Negative (Type II) | 2 | 0.8% | Missed Diagnosis | 2.0% |

**Supplementary Table 1. Detailed Misclassification Analysis**
Error breakdown by diagnostic category. Percentages are relative to each group’s total sample size (N = 70 per MCI/AD group; N = 100 for control group).

Statistical notes: Error rates calculated as proportion of true group misclassified. Percentages rounded to nearest tenth.
