## Supplementary Table S2 for "Linguistic and Acoustic Biomarkers from Simulated Speech Reveal Early Cognitive Impairment Patterns in Alzheimer’s Disease"

**Supplementary Table S2. Summary of Classification Errors and Clinical Implications**

| Model | Total Errors | Error Rate | Dominant Error | Clinical Priority | Recommendation |
| --- | --- | --- | --- | --- | --- |
| AD Classification | 22 | 9.2% | False Negative (77.3%) | Minimize missed AD cases | Lower threshold for AD classification |
| MCI Classification | 29 | 12.1% | False Positive (65.5%) | Balance sensitivity/specificity | Optimize decision boundary |
| Control Classification | 7 | 2.9% | False Positive (71.4%) | Maintain high specificity | Current threshold appropriate |
